# Efficient Generation of Isogenic Primary Human Myeloid Cells using CRISPR-Cas9 Ribonucleoproteins

**DOI:** 10.1101/2020.03.13.991414

**Authors:** Joseph Hiatt, Devin A. Cavero, Michael J. McGregor, David E. Gordon, Weihao Zheng, Jonathan M. Budzik, Theodore L. Roth, Kelsey M. Haas, Ujjwal Rathore, Anke Meyer-Franke, Mohamed S. Bouzidi, Judd F. Hultquist, Jason A. Wojcechowskyj, Krystal A. Fontaine, Satish K. Pillai, Jeffery S. Cox, Joel D. Ernst, Nevan J. Krogan, Alexander Marson

## Abstract

Genome engineering of primary human cells with CRISPR-Cas9 has revolutionized experimental and therapeutic approaches to cell biology, but human myeloid-lineage cells have remained largely genetically intractable. We present a method for delivery of CRISPR-Cas9 ribonucleoprotein (RNP) complexes by nucleofection directly into CD14+ human monocytes purified from peripheral blood, leading to high rates of precise gene knockout. These cells can be efficiently differentiated into monocyte-derived macrophages or dendritic cells. This process yields genetically-edited cells that retain critical markers of both myeloid differentiation and phagocytic function. Genetic ablation of the restriction factor SAMHD1 increased HIV-1 infection more than fifty-fold, demonstrating the power of this system for genotype-phenotype interrogation. This fast, flexible and scalable platform can be used for genetic studies of human myeloid cells in immune signaling, inflammation, cancer immunology, host-pathogen interactions, and beyond, and could facilitate development of novel myeloid cellular therapies.

## Introduction

Myeloid cells are key players in the immune system in health and disease (Germic et al., 2019; Lapenna et al., 2018; Worbs et al., 2017). Monocytes and macrophages function in the immediate arm of the innate immune system, responding to pathogens or tissue damage and helping to regulate and resolve inflammation in tissue. As professional antigen presenting cells, dendritic cells orchestrate the adaptive immune response. Given their central roles, it is unsurprising that myeloid cells have been identified as key players in everything from development and homeostatic regulation to pathogen response, autoinflammatory disease, fibrosis and malignancy (Chao et al., 2020; Engblom et al., 2016; Manthiram et al., 2017; Medzhitov and Janeway, 2000, 1997; Wynn et al., 2013). Improved understanding of the normal and pathogenic behaviors of these cells is crucial to furthering our mechanistic understanding of a broad range of disorders, offering hope for the discovery and advancement of new treatments.

Our ability to identify new therapeutic targets and construct novel cellular interventions has advanced in lockstep with our ability to genetically manipulate relevant primary cell types. For example, mouse genetic approaches have exposed the remarkable diversity of mouse macrophages, and genetic ablation of myeloid subsets paved the way for therapeutic targeting of analogous cells in the clinic (Wynn et al., 2013). CRISPR-Cas9-mediated gene targeting has significantly expanded the potential of once-intractable cell types, facilitating important discovery efforts and enhanced cell therapy approaches in primary T cells (Roth et al., 2018; Schumann et al., 2015; Simeonov and Marson, 2019; Stadtmauer et al., 2020), as well as cures for debilitating genetic diseases using edited hematopoietic stem/progenitor cells (Foss et al., 2019; Wu et al., 2019).

Until now, CRISPR-Cas9 has been inefficient in primary human myeloid cells, limiting functional genetic studies and genome engineering in these key cells of the human immune system. The identification of SAMHD1 as the key restriction factor in myeloid cells that prevents efficient lentiviral transduction (Hrecka et al., 2011; Laguette et al., 2011) led directly to improved approaches for studying innate immunity and has been leveraged to generate more effective dendritic cell vaccines (Norton et al., 2015; Sunseri et al., 2011). Even so, studies of human myeloid cells continue to suffer from the difficulty of accessing and manipulating relevant cell subsets (Lee et al., 2018). Expanding the genetic toolkit with CRISPR-Cas9 would enable further dissection of the genetic circuits underlying the behavior and development of this remarkably diverse class of cells (Geissmann and Mass, 2015; Hancock et al., 2013), offering new insight and more specific, sophisticated targets for therapeutic manipulation.

We report here a robust, flexible, and cost-efficient platform for genetically modifying primary human CD14+ monocytes, which can then be quickly differentiated into monocyte-derived macrophages (MDMs) or monocyte-derived dendritic cells (MoDCs). We demonstrate the utility of using this system to study host-pathogen interactions, however this approach is equally suited to the study of any number of other phenotypic outcomes. The platform is designed to be scalable and is compatible with workflows assessed by microscopy, flow cytometry, and a wide range of other common assays, and is suitable for the interrogation of both cell-intrinsic and non-cell-autonomous behaviors.

## Results

### Efficient Gene Ablation in Primary Myeloid Cells

CD14-positive monocytes are abundant in peripheral blood, representing about 10% of circulating leukocytes (Auffray et al., 2009), and can be differentiated *ex vivo* into monocyte-derived macrophages (MDMs) or monocyte-derived dendritic cells (MoDCs) (Figueroa et al., 2016; Jin and Kruth, 2016), making them the ideal starting point for generation of isogenic primary myeloid cells. Monocytes are isolated from donor blood and immediately subjected to CRISPR-Cas9 ribonucleoprotein (RNP) nucleofection. They are then put into culture, allowing several days for turnover of the targeted gene product under conditions leading to differentiation into MDMs or MoDCs, after which the cells can be subjected to a range of functional, genotypic and phenotypic assays **(Figure 1a)**. A survey of conditions for the Lonza 4D Nucleofector identified pulse code DK-100 in buffer P2 as optimally balancing editing efficiency, cell survival and cell morphology **(Figure S1a)**. Using this approach, we showed robust, guide-sequence-dependent knockout of genes expressed in myeloid cells by immunoblot of the gene product at day 7 of differentiation **(Figure 1b)**. By testing multiple guides, we were able to reproducibly achieve at least a 75% reduction in targeted protein levels relative to untargeted housekeeping control gene products **(Figure 1c)**. Further, we confirmed consistently robust knockout across biological replicates at the genetic level by Tracking of Indels by DEcomposition (TIDE) analysis (Brinkman et al., 2014), showing that guides against *CXCR4* and *CCR5* led to disruption in greater than 90% of alleles **(Figure 1d)**. This protocol led to reproducible knockout when starting with CD14+ monocytes from freshly isolated peripheral blood mononuclear cells (PBMC), from cryopreserved PBMC, or from isolated-then-cryopreserved CD14+ monocytes **(Figure S1b)**, allowing for a flexible workflow and enabling iterative experiments on consistent biological samples. Collectively, these data demonstrate optimized knockout of targeted genes in primary human myeloid cells using CRISPR-Cas9 RNPs.

**Figure 1.**
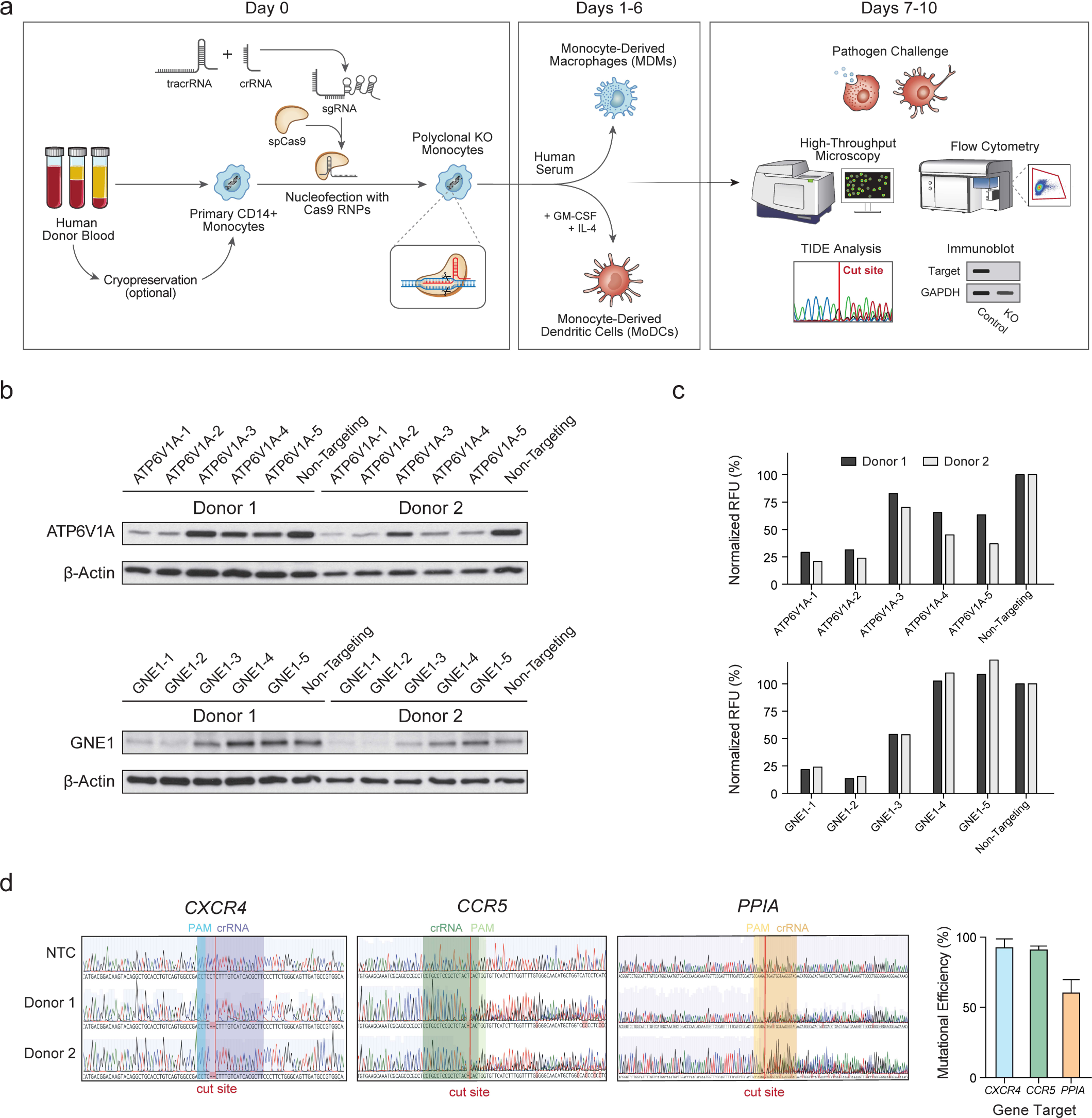
A flexible platform for CRISPR editing of human myeloid-lineage cells. **(a)** A generalized schematic of the platform. Human CD14+ monocytes are isolated from blood by density gradient separation of PBMC followed by magnetic negative selection. Either PBMC or monocytes may be cryopreserved for later editing **(Figure S1b)**. Cells are then nucleofected with preformed CRISPR-Cas9 RNPs and cultured under MDM- or MoDC-generating conditions. After allowing for differentiation and washout of the targeted gene product, cells can be subjected to a wide variety of functional, phenotypic and genotypic studies to assess the knockout efficiency and the function of the targeted gene product. **(b)** Guide-sequence-dependent knockout of targeted genes leads to loss of gene products. CD14+ monocytes were nucleofected with RNPs containing one of five distinct guide sequences against the indicated gene or a scrambled non-targeting control, cultured under MDM-generating conditions, and then lysed for immunoblot analysis. Blots show targeted gene protein product and untargeted housekeeping gene product β-actin protein levels in cells from two blood donors. **(c)** Knockout was quantified by digital densitometry and normalized on a per-sample basis in relative fluorescence units (RFU) to untargeted housekeeping control protein β-actin. **(d)** Genomic analysis of knockout target sites allows for quantification of mutational efficiency independent of gene product expression. Left, representative chromatograms of non-targeting control (top) and edited (middle, Donor 1; bottom, Donor 2) sequences, with crRNA sequence, cut site and protospacer-adjacent motif (PAM) highlighted; right, quantification of editing efficiency by TIDE. Bars represent mean ± SD of two (*PPIA*) or four (*CXCR4, CCR5*) biological replicates.

### Edited CD14+ Monocytes Differentiate Robustly into MoDCs and MDMs

We next sought to establish that these edited cells could differentiate into both MDMs and MoDCs. CD14+ monocytes were isolated, edited at the *CXCR4* locus or left unperturbed, and put into culture under MDM-differentiating conditions in Iscove’s Modified Dulbecco’s Medium (IMDM) with 20% human male AB serum; or under MoDC-differentiating conditions in IMDM with 1% human male AB serum, 50ng/mL IL-4 and 50ng/mL GM-CSF. After six days of culture both edited and unperturbed monocytes had differentiated equally into MDMs, characterized by high levels of expression of CD14, CD16, CD11b and CD206 **(Figure 2a)**. Cells cultured under MoDC-generating conditions displayed upregulation of CD11b, CD11c, and HLA-DR, as expected. We did observe some discrepancies in CD14 and HLA-DR expression between the nucleofected and un-nucleofected control cells, suggesting differences in differentiation or maturation of MoDCs under these conditions **(Figure 2b)**.

**Figure 2.**
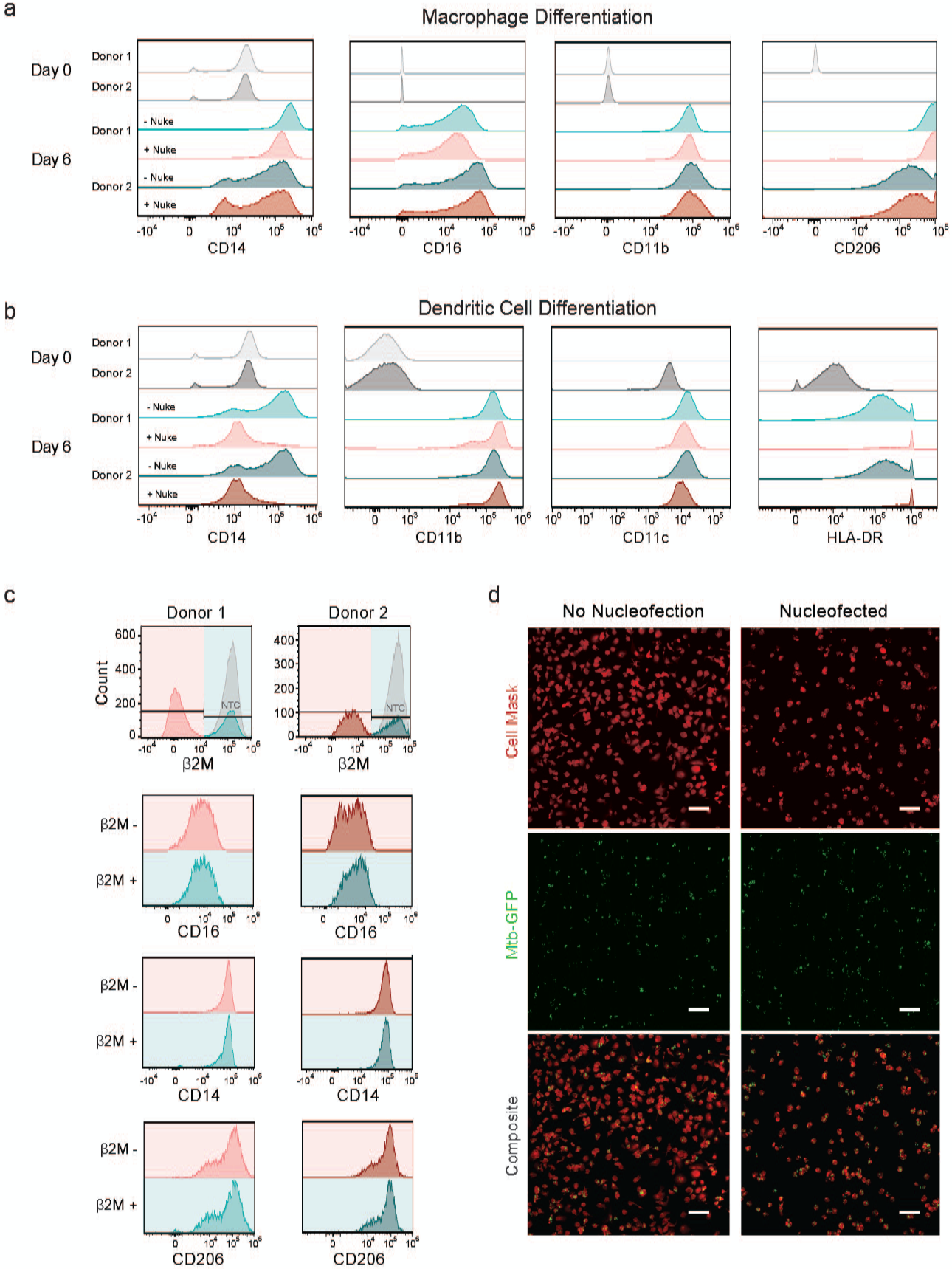
CRISPR-Cas9 mediated gene knockout preserves key aspects of differentiation and function in targeted myeloid cells. (**a)** Immunophenotypic analysis of edited cells is comparable to unedited cells at day 6 of culture under MDM-generating conditions; two representative donors shown. CD206 staining was available for only one donor at day 0. “- Nuke”, un-nucleofected cells; “+ Nuke”, cells nucleofected with *CXCR4*-targeting RNPs. **(b)** Immunophenotypic analysis of edited and unedited cells cultured under MoDC-generating conditions reveals similar CD11b and CD11c expression and differences in CD14 and HLA-DR levels; two representative donors are shown. CD11c and HLA-DR staining were available for only one donor at day 0. **(c)** Among cells subjected to CRISPR-Cas9 RNP nucleofection, immunophenotypic differences were not observed between the cells that bear the desired β2m knockout (pink) and those that do not (teal) after 7 days of MDM differentiation. (**d)** Representative images of unperturbed (left) and RNP-nucleofected (right) MDMs infected with GFP-expressing *M. tuberculosis* (Mtb-GFP) show that CRISPR-Cas9-targeted cells remain competent to phagocytose living pathogens. Top, membrane staining with Cell Mask Far Red; middle, Mtb-GFP; bottom, composite. Scale bars represent 100µm.

To more deeply assess whether CRISPR-Cas9 editing altered the differentiation process, we sought to determine whether editing occurred preferentially in specific immunologic subpopulations. We employed an intermediate-efficiency guide against an easily stained surface antigen, Beta-2-microglobulin (β2m), in order to generate a roughly even mix of β2m-negative and β2m-positive cells that had all been exposed to the same nucleofection and had been cultured together. These cells were subjected to MDM differentiation for seven days before co-staining for β2m and several myeloid phenotypic markers **(Figure 2c)**. The β2m-positive cells thus act as in-well controls for the β2m-negative, knockout cells. We observed no phenotypic difference between the β2m-positive and β2m-negative cells, suggesting that the process of CRISPR gene ablation does not appreciably select for a subset of cells, nor skew MDM differentiation.

We then asked if edited primary myeloid cells not only acquired key markers of differentiation, but also retained critical functions of mature macrophages. We evaluated the ability of edited and unedited MDMs to phagocytose *Mycobacterium tuberculosis*, a deadly pathogen found inside macrophages and dendritic cells during human infection (Wolf et al., 2007). Internalization of *M. tuberculosis*, a key stage in the bacterial lifecycle, is mediated at least in part by phagocytosis by host macrophages (Ernst and Wolf, 2006; Srivastava et al., 2016). When we exposed either unperturbed MDMs or MDMs nucleofected with RNPs targeting the irrelevant *CXCR4* gene, we found that both cell populations were capable of phagocytosing the pathogens at a high rate, leaving few extracellular bacteria **(Figure 2d)**. Edited human myeloid cells can retain phagocytic ability, consistent with successful differentiation despite Cas9 RNP nucleofection, and thus represent an ideal platform for functional assays.

### Functional Testing of Host Factors in Isogenic Primary Myeloid Cells

Finally, we sought to establish that this platform is suitable to test the function of genetic factors that control complex cellular behaviors. We targeted the host viral restriction factor SAMHD1, which blocks lentiviral infection (including HIV-1 infection) by depleting intracellular pools of dNTPs needed for reverse transcription (Gelais and Wu, 2011; Hrecka et al., 2011; Laguette et al., 2011; Sunseri et al., 2011). We employed five distinct guides against *SAMHD1*, or a non-targeting control guide, to attempt to perturb the *SAMHD1* gene directly in primary human monocytes from four different blood donors. After 7 days of differentiation into MDMs, cells were exposed to the CCR5-tropic, chimeric HIV-1 clone LAI-YU2. Two days later, cells were stained with Hoechst dye and probed for the HIV-1 antigen p24, which is indicative of productive infection, then quantified by high-throughput microscopy **(Figure 3a-b, Figure S2a-b)**. SAMHD1 deletion resulted in a significant, greater-than-fifty-fold increase in HIV infection. (Guide SAMHD1-1: 51.74-fold increase in infection relative to non-targeting, 95% confidence interval 14.17-89.32; SAMHD1-2: 50.30-fold increase, 95% CI 12.73-87.88; SAMHD1-3: 39.17-fold increase, 95% CI 1.594-76.74; results of a repeated measures one-way ANOVA with Dunnett’s multiple comparison test.) We observed a clear correlation between knockout efficiency as measured by protein or DNA **(Figure 3c-d)** and the degree of HIV infection, consistent with a causal relationship **(Figure S2c)**. This dramatic effect upon ablation of a viral restriction factor clearly demonstrates that this platform is suitable for the functional assessment of host genes in primary human myeloid-lineage cells.

**Figure 3.**
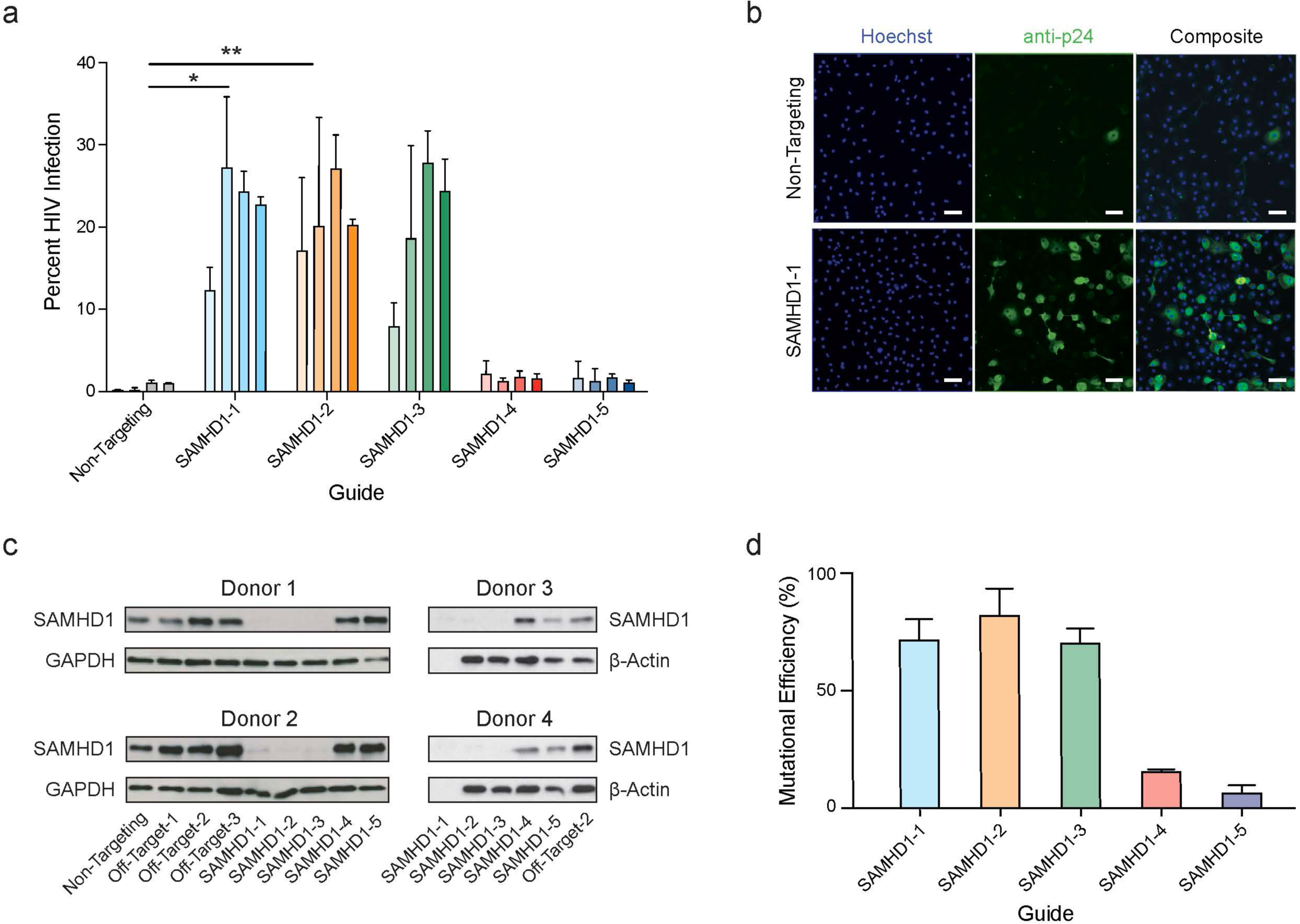
Generation of isogenic monocyte-derived macrophages for functional evaluation of an HIV-1 host factor. **(a)** *SAMHD1*-targeted or non-targeting control MDMs from four independent, HIV-negative blood donors were infected with HIV-1. Plot displays the percentage of cells productively infected after 48-hour exposure. Guides that most efficiently ablated the gene caused statistically significant increases in infection as assessed by one-way ANOVA followed by Dunnett’s test. * = p < 0.05, ** = p < .01. See also **(Figure S2b). (b)** Representative images of HIV-1 infection from Donor 3 comparing cells nucleofected with control non-targeting RNPs (top) to cells nucleofected with RNPs made from guide SAMHD1-1 (bottom). Left, Hoechst; middle, staining of the HIV-1 antigen p24; right, composite. Scale bars represent 100µm. For representative images of all guides see (**Figure S2a**). Quantification of *SAMHD1* knockout by immunoblot **(c)** and sequencing **(d)**. No protein sample was available for guide SAMHD1-1 in donors 3 and 4; Sanger sequencing was analyzed for mutational efficiency by TIDE, bars represent mean ± SD for at least three biological replicates, see also **(Figure S2c)**.

## Discussion

Building on previous work using CRISPR-Cas9 to render primary human immune cells genetically tractable (Hultquist et al., 2019; Hung et al., 2017; Nguyen et al., 2019; Schumann et al., 2015), we have developed a robust, flexible, and high-throughput-compatible platform for the genetic manipulation of primary human myeloid cells. We have optimized cell isolation, culture and CRISPR-Cas9 RNP nucleofection conditions to allow for rapid and inexpensive generation of isogenic modified cells that can then be differentiated and flexibly incorporated into downstream biochemical and phenotypic assays. Knockout can be easily achieved at programmed gene targets, and the resulting cells largely retain critical markers and key functional characteristics of differentiated myeloid cells. Differentiated, edited, monocyte-derived macrophages remain capable of phagocytosis of living pathogens, and we demonstrate that this editing system can be incorporated into existing assays to study complex biological phenotypes such as host-pathogen interactions.

The platform we present here improves significantly upon existing technology to manipulate human myeloid cells. Genetic perturbation to date has rested largely upon RNAi technologies, which compared to CRISPR-based approaches have higher off-target effects and result in transient, incomplete knockdown, rather than knockout (Housden and Perrimon, 2016); or on lentiviral transduction, which is inefficient in these cells due to high expression of SAMHD1 (Gelais and Wu, 2011; Goujon et al., 2007). This has hampered both the mechanistic investigation of these important, functionally diverse cells and the development of strategies to employ these cells as therapeutics. For these reasons, other groups have previously sought workarounds to make myeloid cells genetically tractable. One approach is to edit hematopoietic progenitor/stem cells and differentiate them into myeloid lineage cells (Kang et al., 2015; Lee et al., 2018; Sontag et al., 2017). Comparatively, this route is difficult, expensive, and time-consuming (Sugimura et al., 2017). Another approach is to edit fully differentiated macrophages, which has shown some functional success in the literature (Barkal et al., 2017; Haney et al., 2018), though this also has limitations. The approach is specific to macrophages, rather than allowing for flexible differentiation, and is limited by the lifespan of mature macrophages in culture. As a result, phenotypic assessment has been limited to short-time-scale assays soon after editing, when cells display appropriate levels of genetic editing but may still have variable concentrations of a targeted gene’s protein product remaining in the cell.

This platform for editing primary human monocytes before differentiation ameliorates these previous limitations. Cells may be differentiated into MoDCs or MDMs; knockout is robust, permanent and can be rapidly and inexpensively iterated by substituting guide RNA sequences; there is adequate time for turnover of the targeted gene product and for subsequent multi-day functional and phenotypic assessment; and the cells are generated in only one week. Isolated cells can be cryopreserved before editing, allowing additional experimental flexibility. Furthermore, all of the reagents and equipment are readily available. We anticipate that these features – which have proven key to the success of CRISPR-Cas9 RNP-based editing of T cells – will facilitate a diverse set of research endeavors and hopefully accelerate the development of myeloid cell therapies.

## Acknowledgements

We thank all members of the Marson lab and Krogan lab for support and advice. We are grateful for discussions with B.R. Conklin, M. Ott, D.G. Russel, M. Jost, V. Ramani, G. Alberts, S. Pyle, D. Sainz and G. Ehle. We are grateful for the generosity of our blood donors and the help of Y.D. and the research support team at Vitalant. R. Gummuluru and S. Stanley graciously shared the HIV-1 plasmid and the Mtb-GFP bacteria, respectively. J.H. was supported by the UCSF MSTP (T32GM007618). K.M.H. is supported by the National Science Foundation (1650113). The Marson lab has received gifts from J. Aronov, G. Hoskin, K. Jordan, B. Bakar, the Caufield family and has received funds from the Gladstone Institutes, the Innovative Genomics Institute (IGI) and the Parker Institute for Cancer Immunotherapy (PICI). A.M. holds a Career Award for Medical Scientists from the Burroughs Wellcome Fund, the Lloyd J. Old STAR award from the Cancer Research Institute (CRI) and is an investigator at the Chan Zuckerberg Biohub. The Krogan Laboratory has received research support from Vir Biotechnology and F. Hoffmann-La Roche. The Marson, Krogan and Cox labs have received funding from the BioFulcrum Viral and Infectious Disease Research Program. This work was supported by funding from the James B. Pendleton Charitable Trust and by NIH grants P50 AI150476 (A.M., N.K.), U19 AI135990 (A.M., N.J.K., J.S.C.), P01 AI063302 (N.J.K., J.S.C), R01 AI150449 (S.K.P.), and R01 AI124471 (J.D.E).

## Author Contributions

Conceptualization: JH, DAC, MJM, TLR, MSB, KAF, SKP, NJK, AM. Investigation: JH, DAC, MJM, DEG, WZ, JMB, KMH, UR, AMF, JAW. Resources: AMF, KAF, JSC, JDE, NJK, AM. Formal analysis: JH, DAC, MJM, DEG. Supervision: JH, KAF, JSC, JDE, NJK, AM. Funding acquisition: KAF, JSC, JDE, NJK, AM. Writing – original draft preparation: JH, DAC, MJM, AM. Writing – review and editing: JH, DAC, MJM, WZ, TLR, KMH, MSB, JFH, JAW, KAF, JDE, NJK, AM.

## Declaration of Interests

The authors declare competing financial interests: T.L.R. is a co-founder of Arsenal Biosciences. A.M. is a co-founder of Spotlight Therapeutics and Arsenal Biosciences. A.M. has served as an advisor to Juno Therapeutics, is a member of the scientific advisory board at PACT Pharma, and is an advisor to Trizell. A.M. owns stock in Arsenal Biosciences, Spotlight Therapeutics and PACT Pharma. The Marson lab has received sponsored research support from Juno Therapeutics, Epinomics, Sanofi, and a gift from Gilead. The Krogan Laboratory has received research support from Vir Biotechnology and F. Hoffmann-La Roche.

**Please note inline figures are at slightly reduced resolution. See the end of this document for full-resolution main and supplementary figures.**

## FIGURE LEGENDS

**Figure S1.**
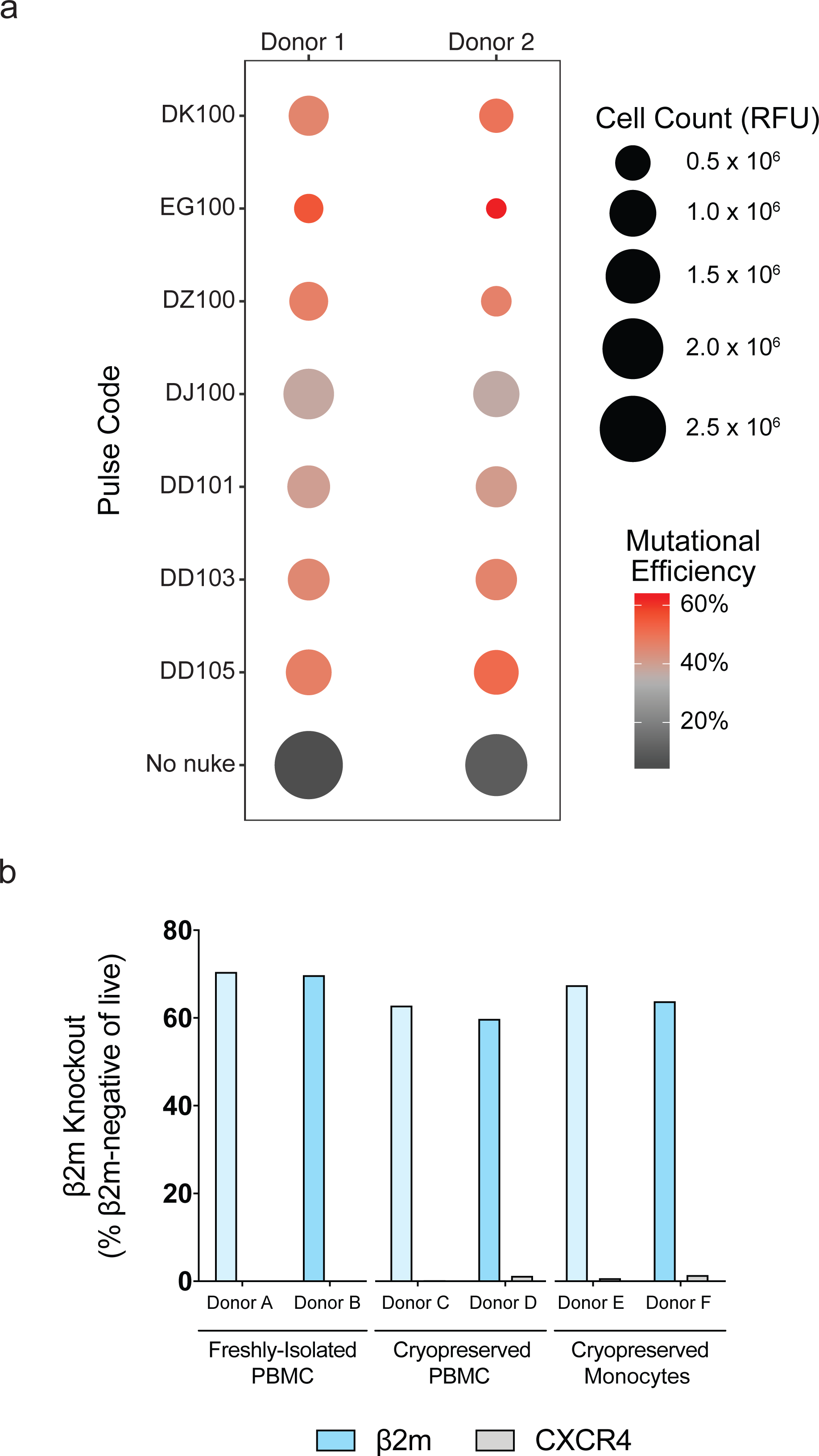
Optimization of knockout in fresh and cryopreserved CD14+ monocytes. (a) Lonza nucleofection. pulse code optimization of *SAMHD1* knockout using guide SAMHD1-2 in CD14+ monocytes freshly isolated from blood of two healthy donors. All nucleofections occurred in buffer P2. Color indicates mutational efficiency determined by TIDE, size of circles indicates relative surviving cell count measured by CellTiter-Glo fluorescent assay in relative fluorescence units (RFU), mean of technical triplicates. Cells were also visually monitored for health and morphology. **(b)** Knockout efficiency in CD14+ monocytes freshly isolated from blood (Donors A and B), CD14+ monocytes freshly isolated from cryopreserved PBMC (Donors C and D), and cryopreserved CD14+ monocytes (Donors E and F). Knockout at the targeted β2m locus was determined by TIDE compared to the non-targeting control (blue bars, left side of each donor); grey bars represent the TIDE knockout efficiency at the β2m locus of off-target *CXCR4* RNPs (grey bars, right side of each donor.)

**Figure S2.**
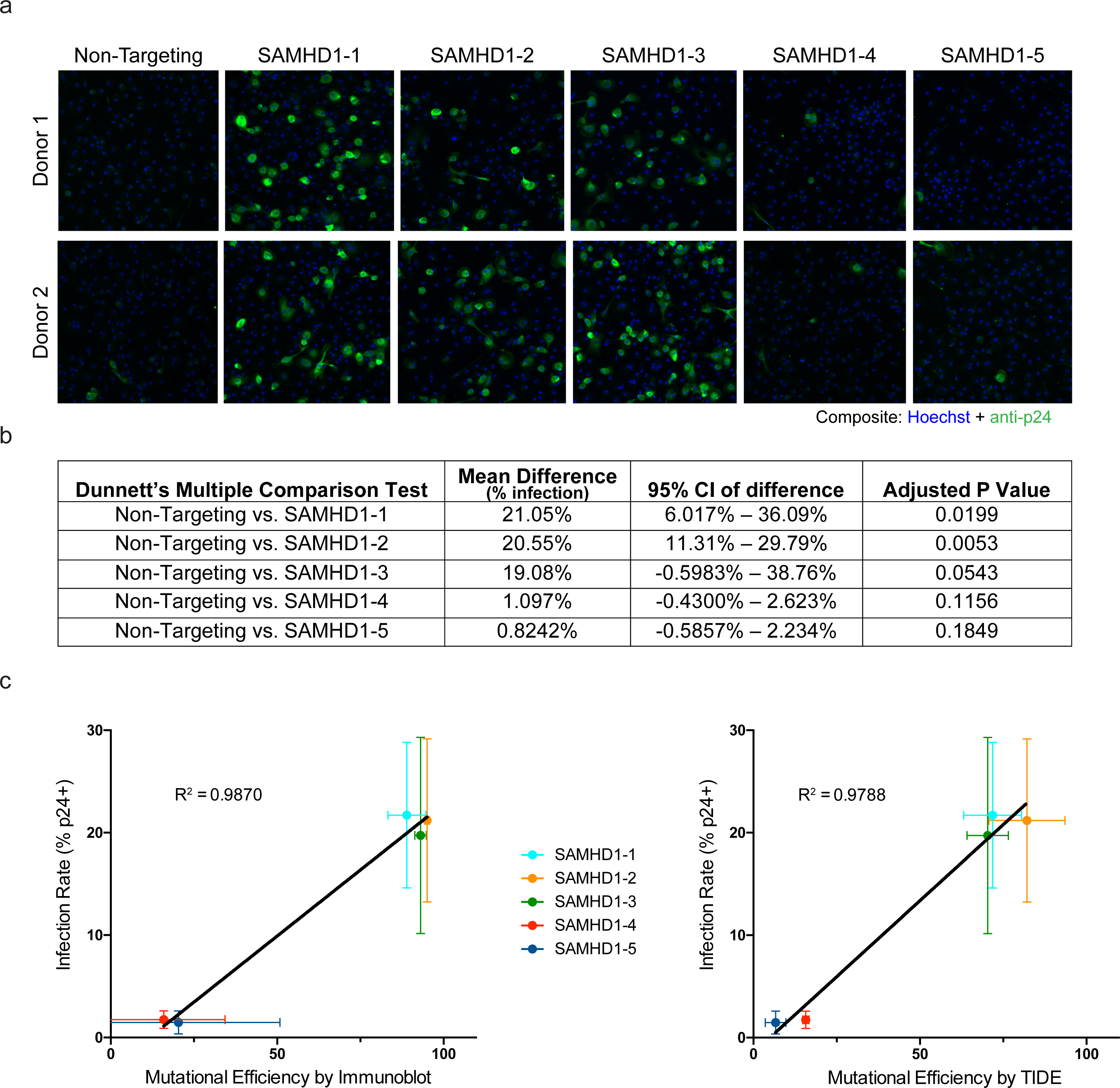
Knockout of the host restriction factor SAMHD1 leads to increases in HIV infection in a manner correlated with guide efficiency. **(a)** Representative composite images of HIV-1 infection from Donors 1 and 2 comparing cells nucleofected with RNPs made from each *SAMHD1*-targeting crRNA and cells nucleofected with control non-targeting RNPs. Blue, Hoechst; green, staining of the HIV-1 antigen p24. **(b)** Statistical comparison of infection rates (percent of cells staining positive for HIV-1 antigen p24). All values were generated by GraphPad Prism using a repeated measures one-way ANOVA followed by Dunnett’s multiple comparison test. **(c)** Correlation of HIV-1 infection rate and knockout efficiency as measured by immunoblot (left panel) or Sanger sequencing quantified by TIDE (right panel). Points represent mean across four donors, except for immunoblot of guide 1 (n = 2) and TIDE of guides 1, 3 and 4 (n = 3); error bars represent standard deviation. Line of best fit and R^2^ values were generated by linear regression in GraphPad Prism.

**Figure S3.**
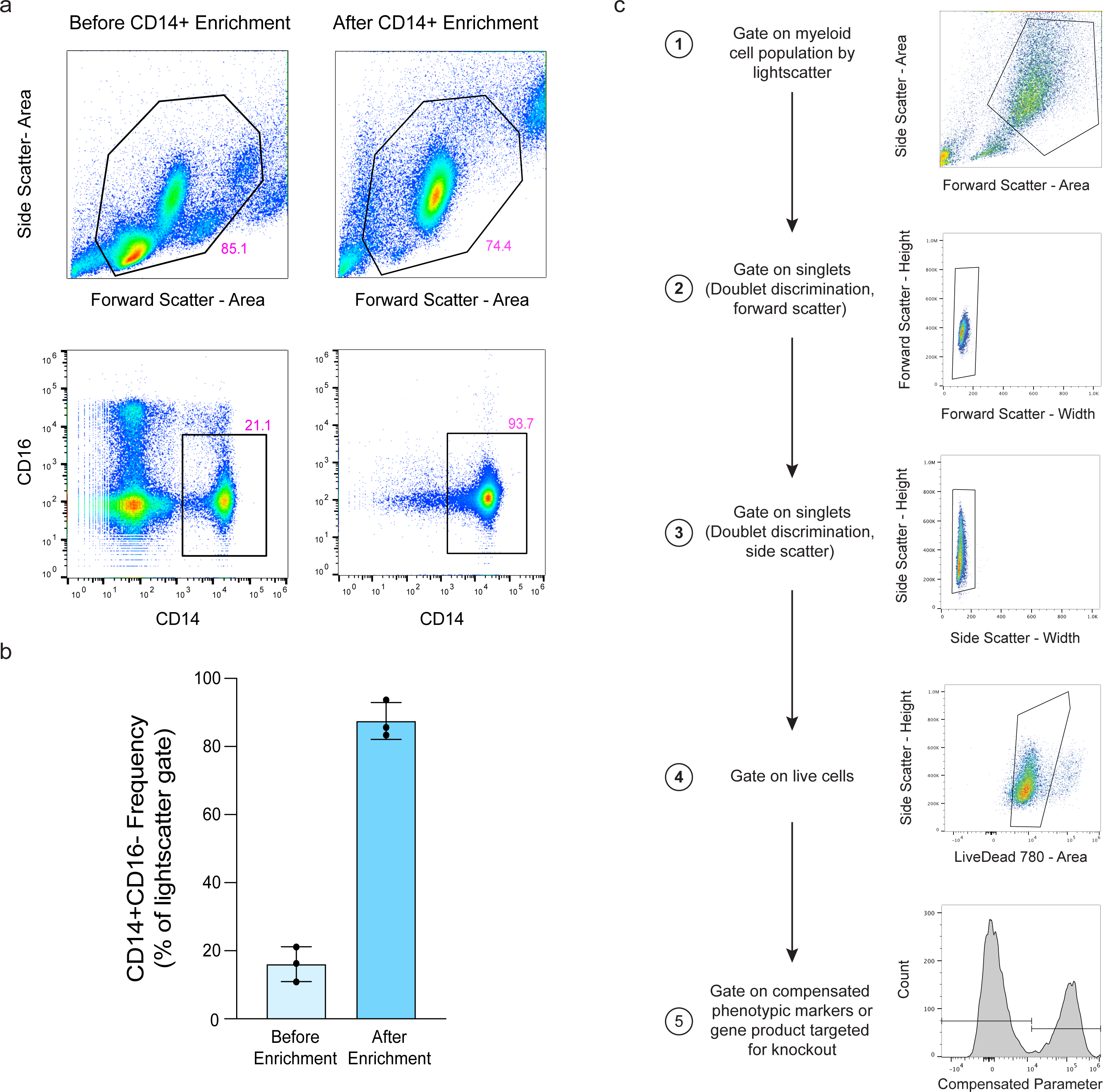
Enrichment of CD14+ cells from PBMC and flow cytometry gating strategy. **(a)** Representative lightscatter (top) and CD14 vs. CD16 staining (bottom) of density-separated PBMC (left) and negatively selected CD14+ monocytes (right). **(b)** Quantification of enrichment by magnetic negative selection across three donors. Bars represent mean ± SD with individual points marked. **(c)** Panels illustrating five-step sample gating strategy for myeloid cells.

## STAR Methods

### KEY RESOURCES TABLE

Please see attached file.

### LEAD CONTACT AND MATERIALS AVAILABILITY

Questions and requests for resources and reagents should be directed to the Lead Contact, Dr. Alexander Marson.

### EXPERIMENTAL MODEL AND SUBJECT DETAILS

Human 293T/17 cells were obtained from the UCSF Cell Culture Facility and were cultured in Dulbecco’s Modified Eagle Medium (DMEM) supplemented with 10% fetal calf serum (FCS, Invitrogen 26140079) at 37°C in 5% CO_2_. Human peripheral blood mononuclear cells (PBMC) were isolated from TRIMA residuals (Vitalant Research Institute) via density-gradient separation according to institutional safety protocols. CD14+ monocytes were isolated from PBMC by magnetic negative selection and cultured at 37°C in 5% CO_2_ on non-treated flat-bottom culture plates in 1X Iscove’s Modified Dulbecco’s Medium (IMDM, Gibco 12440053), supplemented as follows. For dendritic cell differentiation: 1% Human AB Serum (Valley Biomedical HP1022HI), Penicillin-Streptomycin (100IU and 100µg/mL, respectively, Corning 30-002-CI), 1mM Sodium Pyruvate, 50ng/mL GM-CSF (Life Technologies PHC2015), 50ng/mL IL-4 (Life Technologies PHC0045); for macrophage differentiation: 20% Human AB Serum, Penicillin-Streptomycin, 1mM Sodium Pyruvate.

## METHOD DETAILS

### Primary Human Monocyte Isolation and Enrichment

One TRIMA residual (50mL of human peripheral blood enriched in mononuclear cells) from Vitalant Research Institute was processed for each donor using BSL2* precautions in accordance with institutional safety guidelines. Blood was diluted 1:1 with a 4mM EDTA PBS solution, then layered on top of Ficoll-Paque Plus density gradient medium (Sigma GE17-1440-03) in SepMate tubes (StemCell Technologies 85450). After centrifugation at 1200xg for 10 minutes at room temperature, blood separated into plasma, PBMC, granulocyte, and erythrocyte layers. The plasma was aspirated allowing the PBMC buffy coat to be decanted. It was then suspended in 2mM EDTA PBS and centrifuged at 400xg for 10 minutes at room temperature. PBMC were washed twice with 2mM EDTA PBS by centrifugation for 10 minutes at 300xg and then again at 200xg. Cells were counted, pelleted a final time at 200xg for 10 minutes, and resuspended at a concentration of 50 × 10^6^ cells/mL in MACS buffer (PBS, 2mM EDTA, 0.5% BSA). At this point, 0.5 × 10^6^ PBMC from each donor were removed for quality control flow cytometric analysis as described below. To enrich for CD14+ monocytes, 50µL/mL of CD14+ negative selection cocktail (StemCell 19359) was added and incubated for 5 minutes at room temperature. After incubation, suspensions were placed in an EasyEights EasySep Magnet (StemCell 18103) tube rack for 5 minutes. Enriched monocytes were carefully pipetted from the magnet, and 0.5 × 10^6^ monocytes from each donor were again removed for flow cytometric quality control analysis as described below (See also **Figure S3a-b)**.

Where noted, isolated PBMC or CD14+ monocytes were cryopreserved in FBS with 10% DMSO in a Mr. Frosty Freezing container (ThermoFisher 5100) stored at −80°C for 1-4 days before transfer to liquid nitrogen. Cryopreserved PBMC were carefully thawed and washed twice with MACS buffer before incubation with negative selection cocktail as described above.

### Formation of Cas9-Ribonucleoproteins

Cas9 ribonucleoproteins were produced as previously described (Hultquist et al., 2019). Briefly, lyophilized crRNA and tracrRNA (Dharmacon, see Key Resources Table Oligonucleotides) were resuspended at a concentration of 160µM in 10mM Tris-HCL (7.4 pH) with 150mM KCl and immediately used or frozen at −80°C. No more than one thaw was allowed for any RNP reagent. Equal volumes of crRNA and tracrRNA were incubated at 37°C for 30 minutes to form an 80 µM guide RNA duplex; this was then incubated with an equal volume (2:1 RNA:Cas9 molar ratio) of 40µM Cas9 protein (UC Macrolab) at 37°C for 15 minutes to form RNPs at 20µM. RNPs were used immediately or stored at −80°C.

### Nucleofection of Cas9-Ribonucleoproteins into Primary Human Monocytes

Isolated CD14+ monocytes were counted and 0.5-1 × 10^6^ cells per nucleofection reaction were spun down at 200xg for 8 min. Supernatant was carefully and completely aspirated, and cells were resuspended in 20µL/reaction of room-temperature Lonza nucleofection buffer P2 (Lonza V4XP-2024). The cell suspension was gently mixed with 2.5µL/reaction of appropriate RNP and then pipetted into a 96-well-format nucleofection cuvette for the Lonza 4D X unit or Shuttle unit (Lonza). Except where explicitly stated, cassettes were nucleofected with code DK-100, immediately supplemented with 80µL pre-warmed culture medium, and rested in a dark, 37°C, 5% CO_2_ incubator for 15-30 minutes. Subsequently cells were moved to a prepared non-treated, flat-bottom culture plate pre-filled with appropriate media for differentiation and subsequent analysis. One nucleofection reaction of 0.5 × 10^6^ cells is sufficient to seed three wells of a 96-well plate or one well of a 48-well plate.

### *In vitro* Differentiation of Monocyte-Derived Macrophages and Dendritic Cells

Cells were cultured in flat-bottom, non-treated cell culture plates in either 96-well (Corning 351172) or 48-well (Corning 351178) format. Twenty-four hours after nucleofection, after visually confirming cell adherence, the entire volume of media was exchanged for fresh, pre-warmed culture media. Three and five days after nucleofection, half of the culture media was removed and replaced with fresh media pre-warmed to 37°C. Media formulations are as follows. For MoDCs: 1X IMDM (Gibco 12440053), 1% Human AB Serum (Valley Biomedical HP1022HI), Penicillin-Streptomycin (100IU and 100µg/mL, respectively, Corning 30-002-CI), 1mM Sodium Pyruvate, 50ng/mL GM-CSF (Life Technologies PHC2015), 50ng/mL IL-4 (Life Technologies PHC0045); for MDMs: 1X IMDM, 20% Human AB Serum, Penicillin-Streptomycin, 1mM Sodium Pyruvate.

### Flow Cytometric Staining and Analysis

To detach adherent MDMs and MoDCs for downstream flow cytometric analysis, cells were first spun for 5 minutes at 300xg, media was removed, and cells were incubated in Accutase Cell Detachment Solution (ThermoFisher Scientific 00-4555-56) for 15 minutes at 37 ° C. Cells were then transferred to V-bottom 96-well plates and resuspended in MACS buffer (PBS, 2mM EDTA, 0.1% Bovine Serum Albumin). Monocytes and PBMC did not require lifting.

To assess the quality of the CD14+ negative selection, samples before and after CD14 enrichment were stained with antibodies against CD14-PE (1:25)(Miltenyi 130-110-519) and CD16-APC (1:25)(Miltenyi 130-106-705).

For phenotypic analysis of MDMs and MoDCs, cells were blocked with Human TruStain FcX (Biolegend 422302) and stained at 4°C in a final volume of 50µL with the following antibodies: CD14-FITC (1:50)(Biolegend 301803), CD16-PE (1:50)(Biolegend 3G8), CD14-PE (1:25)(Miltenyi REA599), CD16-APC (1:25)(Miltenyi REA423), β2-Microglobulin-PE (1:100)(BD 551337), CD11b-BV650 (1:50)(BD 740566), CD11c-PerCP-Cy5.5 (BD 565227), CD206-BV421 (1:50)(BD 566281), LiveDead 510 (1:500)(Tonbo 13-0870-T100), HLA-DR-Pacific Blue (Life Technologies MHLDR28) and GhostDye Red 780 (1:500)(Tonbo 13-0865-T100). Compensation was performed using single-stained UltraComp eBeads Compensation Beads (Life Technologies 01-2222-42) and a mixture of live and killed MDMs or MoDCs for GhostDye Red 780 compensation. Samples were acquired on the Attune NxT Flow Cytometer (ThermoFisher) and analyzed using FlowJo software (FlowJo, LLC). For sample enrichment stains and gating strategy please see **(Figure S3c)**.

### In vitro infection of MDMs by *Mycobacterium tuberculosis*

*M. tuberculosis* was grown to log phase in 7H9 liquid media (BD 271310) supplemented with Middlebrook OADC (Sigma M0678), 0.5% glycerol, 0.05% Tween-80 in roller bottles at 37 ° C. *M. tuberculosis* Erdman strain expressing eGFP under control of the MOP promoter was a gift from Dr. Sarah Stanley’s laboratory. Macrophages were infected with fluorescent *M. tuberculosis* using a modified version of the spinfection protocol as previously described (Watson et al., 2015). Mycobacteria were washed in PBS three times and directly inoculated into the macrophage tissue culture wells at an MOI of 10. Following centrifugation, infected cells were incubated at 37°C for one day post-infection. The wells were washed with PBS and fixed with 4% PFA before staining for microscopy.

### Quantification of *Mycobacterium tuberculosis* Infection

After fixation in 4% PFA and transfer to PBS, plates were stored at 4°C for staining. Cells were permeabilized with 0.1% Triton-X100 (Sigma-Aldrich T8787) for 15 minutes, washed three times with 1X PBS and stained in a volume of 100μL with CellMask Deep Red Stain (1:100,000) (ThermoFisher, H32721) for 30 minutes at room temperature in the dark. After staining, cells were washed three times with 1X PBS and stored in 100µL/well 1X PBS. Cells were then imaged using a Cellomics Arrayscan with a 10X objective (ThermoFisher) and analyzed using the HCS Studio quantitative analysis software (ThermoFisher) by defining cellular events based on the non-specific membrane Deep Red CellMask stain in the 650 channel and then quantifying infection by measuring mean fluorescent intensity in the 488nm channel.

### HIV Production

Macrophage-tropic HIV virus was generated using the HIV LAI-YU2 chimeric molecular clone, from the lab of Rahm Gummuluru. Virus plasmid (12µg) and 100µl PolyJet (Signagen) were diluted separately in two tubes of 625µL serum-free DMEM, then the two solutions were combined and vortexed to mix. After 15 minutes at room temperature, the transfection complexes were added to T175 flasks containing 293T/17 cells, which were gently rocked. Virus-containing culture supernatant was harvested 48 hours post transfection, spun at 400xg for 5 minutes and filtered through a 0.45µm filter. Virus was precipitated by addition of NaCl and 50% PEG-6000 to final concentrations of 300mM and 8.4%, respectively, followed by incubation for two hours at 4°C. Precipitated virus was pelleted for 40 minutes at 3500 rpm, resuspended in complete RPMI media at 50X concentration, frozen in aliquots on dry ice, and stored at −80°C. Virus was titered on wild-type MDMs prior to testing on knockout MDMs.

### HIV Infection, p24 Staining and Imaging

HIV LAI-YU2 was added to macrophages in 96-well format and incubated for 48 hours to allow for infection. After 48 hours, cells were washed twice in PBS (pH 7.4) and fixed in room temperature 4% formaldehyde for 30-60 minutes. Fixative was washed away with two PBS washes, with a final quench in PBS + 2% fetal calf serum (FCS, Invitrogen 26140079). Cells were permeabilized for 5 minutes with saponin buffer (PBS + 0.2% Saponin (Sigma S7900) + 2% FCS). Anti-p24 antibody (AIDS reagent 183-H12-5C) was diluted 1:500 in saponin buffer, 80μL was added to each well, and the plate was incubated overnight at 4°C. Primary antibody was removed and plates were washed 3 times with saponin buffer. Anti-mouse Alexa Fluor 488-conjuated antibody (Invitrogen A-21202) was diluted in saponin buffer (1:500), 80μL was added to each well, and the plate was incubated for 2-3 hours at room temperature protected from light. Secondary antibody was removed, and the plate was washed twice with saponin buffer and once with PBS. Hoescht 33258 (Sigma 861405) was diluted into PBS for a final concentration of 1μg/mL, and 80μL was added to each well followed by a 5 minute incubation at room temperature. Hoescht buffer was removed and the plate was rinsed twice in PBS, then imaged on a CellInsight automated microscope using a 10X objective (ThermoFisher). Cells were identified using nuclear stain, enlarged cellular masks were drawn around the nuclear masks, and p24-positive cells identified by their high average fluorescence in the 488 channel.

### Immunoblotting and Protein Quantification

To prepare protein samples, differentiated myeloid cells were harvested by aspirating the appropriate growth/differentiation media and then adding 100μL of 2.5x reducing sample buffer (RSB, 1.872 mL 0.5M Tris-HCl pH 6.8, 6 mL 50% Glycerol, 3mL 10% SDS, 250μL β-mercaptoethanol, 378 μL 1% bromophenol blue, 1X PBS) directly to each sample well. Cells were lysed by incubating for at least 3 minutes at room temperature, then lysates were transferred to 96-well PCR plates (USA Scientific 1402-9598). Plates were then heated at 95°C for 30 minutes and stored at −20°C. To prepare immunoblots, samples were thawed at room temperature, and 15μL/lane was loaded into an 18-well 4–20% Criterion TGX Gel (Bio-Rad 567-1094). Gels were run at 90 volts for 30 minutes followed by 150 volts for 50 minutes. The samples were then transferred at 0.25 A for 1 hour to a PVDF Membrane (Bio-Rad 1620177). Following protein transfer, membranes were blocked in 4% Milk PBST for 1 hour at room temperature, and then incubated in blocking solution overnight at 4°C with the following antibodies: rabbit monoclonal anti-ATP6V1A (1:1000)(Abcam EPR19270), rabbit polyclonal anti-GNE (1:1000)(Proteintech 25079-1-AP), rabbit polyclonal anti-SAMHD1 (1:1000)(Proteintech 12586-1-AP), rabbit monoclonal anti-β-actin (1:5000)(CST 4970P), mouse monoclonal α-GAPDH (1:5000)(Sigma G8795). Membranes were then washed three times in PBST for 5 minutes each, and then incubated with appropriate secondary antibody for 1 hour at room temperature. Membranes were washed an additional three times, then stained with Pierce ECL Western Blotting Substrate (ThermoFisher 32106). Exposures of the blots were taken with autoradiography film (Thomas Scientific XC59X) and developed with a medical film processor (Konica Minolta Medical & Graphic SRX-101A). Film was scanned at 300 pixels/inch and stored as 8-bit grayscale TIFF files. The level of protein expression for individual samples was quantified in FIJI (Schindelin et al., 2012) by inverting the images, subtracting the background, and determining the fluorescent intensity by measuring the integrated density of individual bands. The protein expression level was then reported as the relative fluorescence of the protein of interest with respect to the paired loading control.

### Quantification of Mutational Efficiency by TIDE Analysis

Cells were lysed in plate format in 50µL QuickExtract DNA Extraction Solution (Lucigen QE09050). Crude lysate was then incubated at 65°C for 20 minutes and 95°C for 20 minutes. Primers were designed with Primer3. PCR amplification was performed using Phusion 2X Master Mix HotStart Flex (New England Biolabs M0536L), 10µM primer pair (see Key Resources Table Oligonucleotides), and approximately 100ng template DNA. PCR amplicons were subsequently sent for cleanup and Sanger sequencing. Mutational efficiency was then determined by comparison of non-targeting and gene-targeting sample chromatograms using the TIDE Web Tool (Brinkman et al., 2014).

### Analysis of Cell Survival by Luminescence

Relative cell viability was determined with CellTiter-Glo (Promega G7570) according to manufacturer’s instructions. Briefly, fresh aliquots of CellTiter-Glo buffer and substrate were mixed and 100μL of the resulting reagent was added to each well of a 96-well culture plate and the plate was put on a shaker for 2 minutes. Lysates were then moved to an opaque-walled 96-well plate (Costar 3912) and incubated at room temperature for 10 minutes. Luminescence was then recorded on an Enspire multimode plate reader (PerkinElmer).

## QUANTIFICATION AND STATISTICAL ANALYSIS

Flow cytometry data was analyzed using FlowJo software. Microscopy data were analyzed using HCS Studio quantitative analysis software. Immunoblots were analyzed with ImageJ. Data were visualized with GraphPad Prism or RStudio. Infection of *SAMHD1*-targeted MDMs was assessed by a repeated measures one-way ANOVA followed by Dunnett’s multiple comparisons test. All statistical details are present in relevant figure legends.

## DATA AND CODE AVAILABILITY

Code was generated only for formatting and visualization of data; all code is available upon request.

